# Investigating causality in associations between education and smoking: A two-sample Mendelian randomization study

**DOI:** 10.1101/184218

**Authors:** Suzanne H. Gage, Jack Bowden, George Davey Smith, Marcus R. Munafo

## Abstract

**Background:** Lower educational attainment is associated with increased rates of smoking, but ascertaining causality is challenging. We used two-sample Mendelian randomization (MR) analyses of summary statistics to examine whether educational attainment is causally related to smoking.

**Methods and Findings:** We used summary statistics from genome-wide association studies of educational attainment and a range of smoking phenotypes (smoking initiation, cigarettes per day, cotinine levels and smoking cessation). Various complementary MR techniques (inverse-variance weighted regression, MR Egger, weighted-median regression) were used to test the robustness of our results. We found broadly consistent evidence across these techniques that higher educational attainment leads to reduced likelihood of smoking initiation, reduced heaviness of smoking among smokers (as measured via self-report and cotinine levels), and greater likelihood of smoking cessation among smokers.

**Conclusions:** Our findings indicate a causal association between low educational attainment and increased risk of smoking, and may explain the observational associations between educational attainment and adverse health outcomes such as risk of coronary heart disease.

## Introduction

Smoking prevalence has been consistently declining in most high income countries over the past 50 years. However, this decline has been seen predominantly in those of higher socio-economic status, which has resulted in an increase in the social patterning of smoking behaviours and widening health inequality (1, 2). Observational studies consistently show that poor educational outcomes are associated with increased smoking (3, 4), but disentangling causality is challenging due to the higher levels of smoking among those from more disadvantaged backgrounds. A recent study found lower educational attainment at age 11 predicted later tobacco use, but that the opposite was true for alcohol and cannabis use (5). Mendelian randomization (MR) is a technique that utilizes genetic variants as unconfounded proxies for an exposure of interest, as a method of ascertaining better evidence for causality than more traditional observational epidemiological methods (6–8).Relatedly, a Mendelian randomization study found that genetic variants that predict educational attainment were associated with a decreased risk of coronary artery disease, indicating a causal relationship between education and the disease (9). One possible pathway through which educational attainment could impact on health is through health behaviours such as smoking.

Here we used two-sample Mendelian randomization (10) to assess the causal relationship between educational attainment and various smoking behaviours, using publicly-available summary statistics from genomewide association studies (GWAS) of educational attainment and smoking behaviours. Specifically, we investigated smoking initiation, heaviness of smoking using self-reported (cigarettes per day) and biomarker (cotinine levels) phenotypes, and smoking cessation. Critical to MR is the assumption that the association of genetic variants acting as a proxy for the exposure operates (directly or indirectly) through the exposure of interest. This assumption can be investigated via the use of a number of complementary MR techniques that rely on orthogonal assumptions (11). We therefore used a range of MR methods as sensitivity analyses to test the robustness of our conclusions.

## Methods

### Education variants

Single nucleotide polymorphisms (SNPs) associated with educational attainment were identified from genomewide significant hits from a GWAS of educational attainment in 305.72 individuals, including 111,349 individuals from the UK Biobank study (12). The GWAS identified 74 SNPs at genomewide significance predicting years in education. Beta coefficients and standard errors were extracted from publicly available summary statistics on 405.72 individuals (the discovery and replication sample, minus data from 23andme). Not all the SNPs were present in the outcome (i.e., smoking) GWAS. Where possible, proxy SNPs (correlated R ≤ 0.9) were identified. The SNPs, proxies, and the analyses they were included in, are detailed in Supplementary Table 1.

### Smoking phenotypes

Smoking initiation was assessed by the Tobacco and Genetics (TAG) consortium (13), as a binary ever/never measure ascertained in 74,053 individuals. Of the 74 possible SNPs associated with years in education, 32 were present in the GWAS of smoking initiation. A further 25 were identified using SNIPA (http://snipa.helmholtz-muenchen.de/snipa/), giving a total of 57 SNPs for these analyses.

Heaviness of smoking measured as self-reported cigarettes smoked per day was also assessed by the TAG consortium, in 38,181 daily smokers. The same 57 SNPs were available for this phenotype as for the initiation phenotype. Heaviness of smoking measured by cotinine levels was assessed by the Cotinine Consortium GWAS (14) in 4,548 daily smokers. Of the 74 possible SNPs associated with education, 72 were present in the cotinine GWAS.

Smoking cessation was assessed in the TAG consortium, in 41,278 individuals who were either current or former smokers. The same 57 SNP were available for this phenotype as for the initiation and heaviness of smoking (cigarettes per day) phenotypes.

### Procedure

Beta coefficients and standard errors of the SNPs associated with years in education were recorded. These SNPs were then identified in the GWASs of the smoking phenotypes, and corresponding log odds ratios or beta coefficients (as appropriate) and standard errors were recoded. The SNP-exposure and SNP-outcome associations for each smoking phenotypes were combined in a fixed effect meta-analysis using inverse-variance weighting (IVW). We also ran a number of sensitivity analyses that provide causal estimates under less stringent assumptions than the traditional MR approach.

First, we implemented MR Egger regression, which relaxes the assumption that the effects of the variants on the outcome are entirely mediated via the exposure [12]. MR-Egger allows for each variant to exhibit some pleiotropy, but assumes that each gene’s association with the exposure is independent in magnitude from its pleiotropic effects (the InSIDE assumption) (15). This is achieved by allowing an intercept term in the weighted regression analysis. The value of the intercept provides an estimate of the degree of pleiotropy affecting the result, while the beta (slope) coefficient represents the causal effect between exposure and outcome adjusted for pleiotropy. We also ran MR Egger analyses using Simulation Extrapolation (SIMEX) correction, which corrects the standard MR-Egger regression coefficients for regression dilution due to uncertainty in the gene-exposure association estimates (16).

Second, we conducted weighted median regression analyses, which can provide a consistent estimate for the true causal effect when up to half of the weight in the MR analysis pertains from genetic variants that exert pleiotropic effects on the outcome (17). We also conducted weighted modal regression analyses, which relax instrumental variable assumptions (18). Finally, we calculated the Cook’s distance and studentized residual measures for the IVW and MR Egger approaches in order to ascertain whether any individual SNPs were outliers, or were too influential in driving the analysis results.

## Results

### Smoking Initiation

Using inverse variance weighted regression, more years in education was strongly associated with reduced likelihood of initiating smoking (beta -0.54, 95% CI -0.71, -0.36). Similarly, using both weighted median MR and weighted modal MR there was strong evidence of an association, although confidence intervals were wider (weighted median beta -0.72, 95% CI -1.12, -0.32). The I^2^_GX_ statistic for these data (which quantifies the expected dilution of the MR-Egger estimate (16)) was 0.62, and so a SIMEX correction was deemed necessary. However, using this approach, the point estimate was directionally opposite (SIMEX-corrected beta 0.44, 95% CI -0.54, 1.42). The intercept indicated weak evidence of pleiotropy (beta -0.01, 95% CI -0.03, 0.00). These results are shown in Table 1 and Figure 1.

**Table 1:** Estimates from various Mendelian randomization methods for the association between education and smoking initiation.

| Method | Beta coefficient | 95% CI | P-value |
| --- | --- | --- | --- |
| IVW MR | -0.538 | -0.713, -0.362 | 9.28x10<sup>-8</sup> |
| Weighted median MR | -0.718 | -1.094, -0.347 | 0.0002 |
| Weighted modal MR | -0.648 | -0.989, -0.307 | 0.0002 |
| MR Egger estimate | 0.320 | -0.344, 0.983 | 0.338 |
| MR Egger intercept | -0.012 | -0.021, -0.003 | 0.010 |
| Egger SIMEX estimate | 0.440 | -0.537, 1.417 | 0.381 |
| Egger SIMEX intercept | -0.014 | -0.027, -0.001 | 0.044 |

Analysis of 74,053 individuals (SNP N=57 out of a possible 74). SNP: Single nucleotide polymorphism; IVW: Inverse variance weighted; MR: Mendelian randomization; SIMEX: Simulation Extrapolation method.

**Figure 1:**
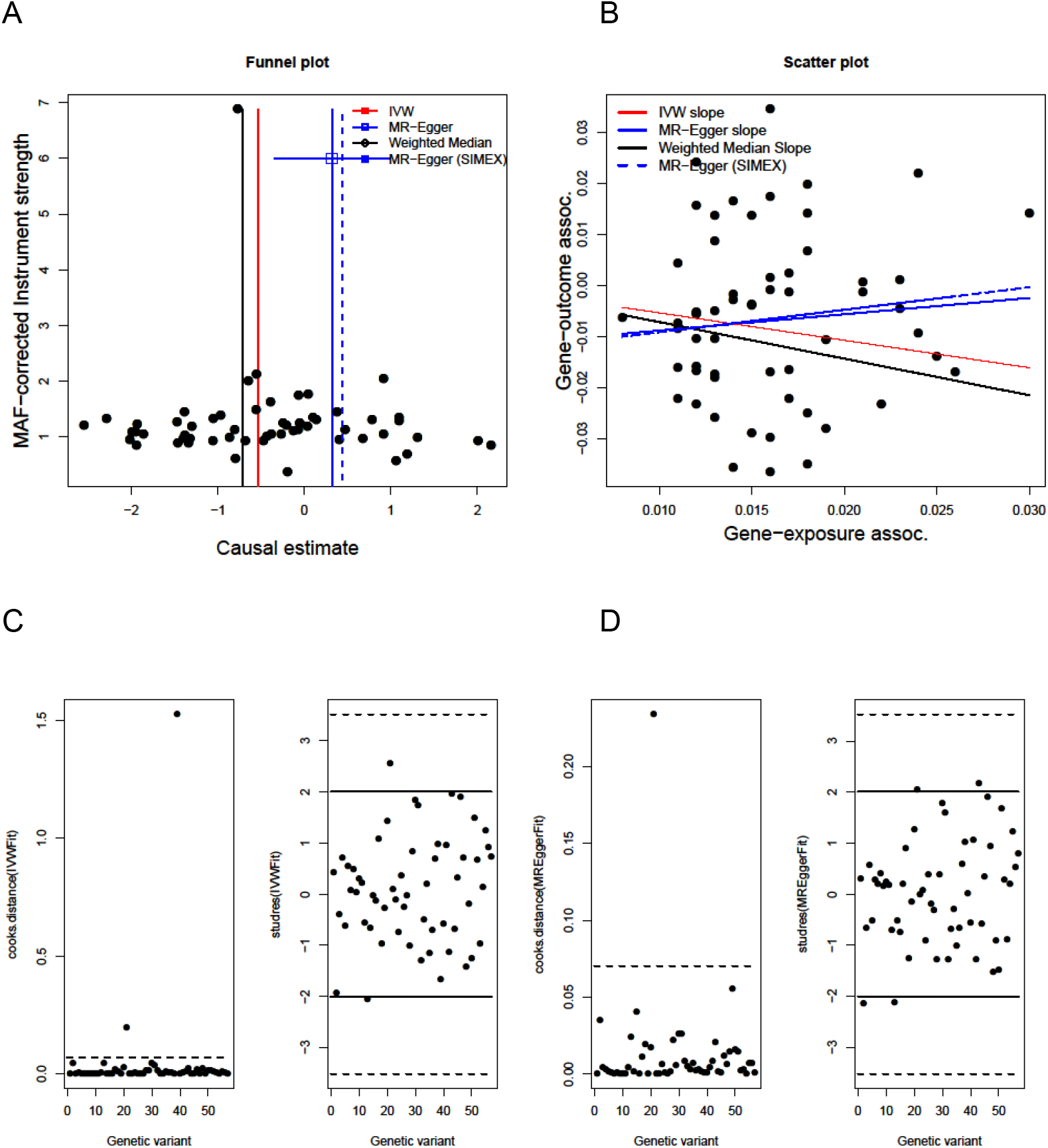
Genetic associations of education with likelihood for smoking initiation. Analysis of 74,053 individuals (SNP N=57 of a possible 74). A) Funnel plot showing minor allele frequency and causal estimate correlations for each SNP. Coloured lines represent the effect sizes of the different regression analyses. B) Scatter plot showing the correlation of genetic associations of education with genetic associations with smoking initiation. Coloured lines represent the slopes of the different regression analyses. C) Cook’s distances and studentized residuals from inverse variance weighted method. D) as C) from MR Egger method, in order to ascertain the presence of outliers.

### Heaviness of Smoking

There was evidence of an association between more years of education and fewer cigarettes smoked per day (beta -2.25, 95% CI -3.81, -0.70). I^2^_GX_ for these analyses was also 0.62. As with the smoking initiation analysis, results were similar when using weighted median MR, although confidence intervals were again wider (beta -2.17, 95% CI -4.40, 0.06), while results using MR Egger were attenuated (SIMEX corrected beta -1.05, 95% CI - 9.42, 7.33). The intercept did not indicate any evidence of pleiotropy (SIMEX corrected beta -0.02, 95% CI -0.15, 0.11). The weighted modal estimate was broadly similar to the IVW and weighted median approach (beta -1.70, 95% CI -5.28, 1.89). These results are shown in Table 2 and Figure 2.

**Table 2:** Estimates from various Mendelian randomization methods for the association between education and cigarettes per day.

| Method | Beta coefficient | 95% CI | P-value |
| --- | --- | --- | --- |
| IVW MR | -2.253 | -3.810, -0.696 | 0.005 |
| Weighted median MR | -2.167 | -4.395, 0.061 | 0.057 |
| Weighted modal MR | -1.698 | -5.284, 1.887 | 0.353 |
| MR Egger estimate | -0.711 | -6.711, 5.289 | 0.813 |
| MR Egger intercept | -0.025 | -0.121, 0.070 | 0.596 |
| Egger SIMEX estimate | -1.045 | -9.423, 7.333 | 0.804 |
| Egger SIMEX intercept | -0.020 | -0.149, 0.109 | 0.759 |

Analysis of 38,181 daily smokers (SNP N=57 out of a possible 74). SNP: Single nucleotide polymorphism; IVW: Inverse variance weighted; MR: Mendelian randomization; SIMEX: Simulation Extrapolation method.

**Figure 2:**
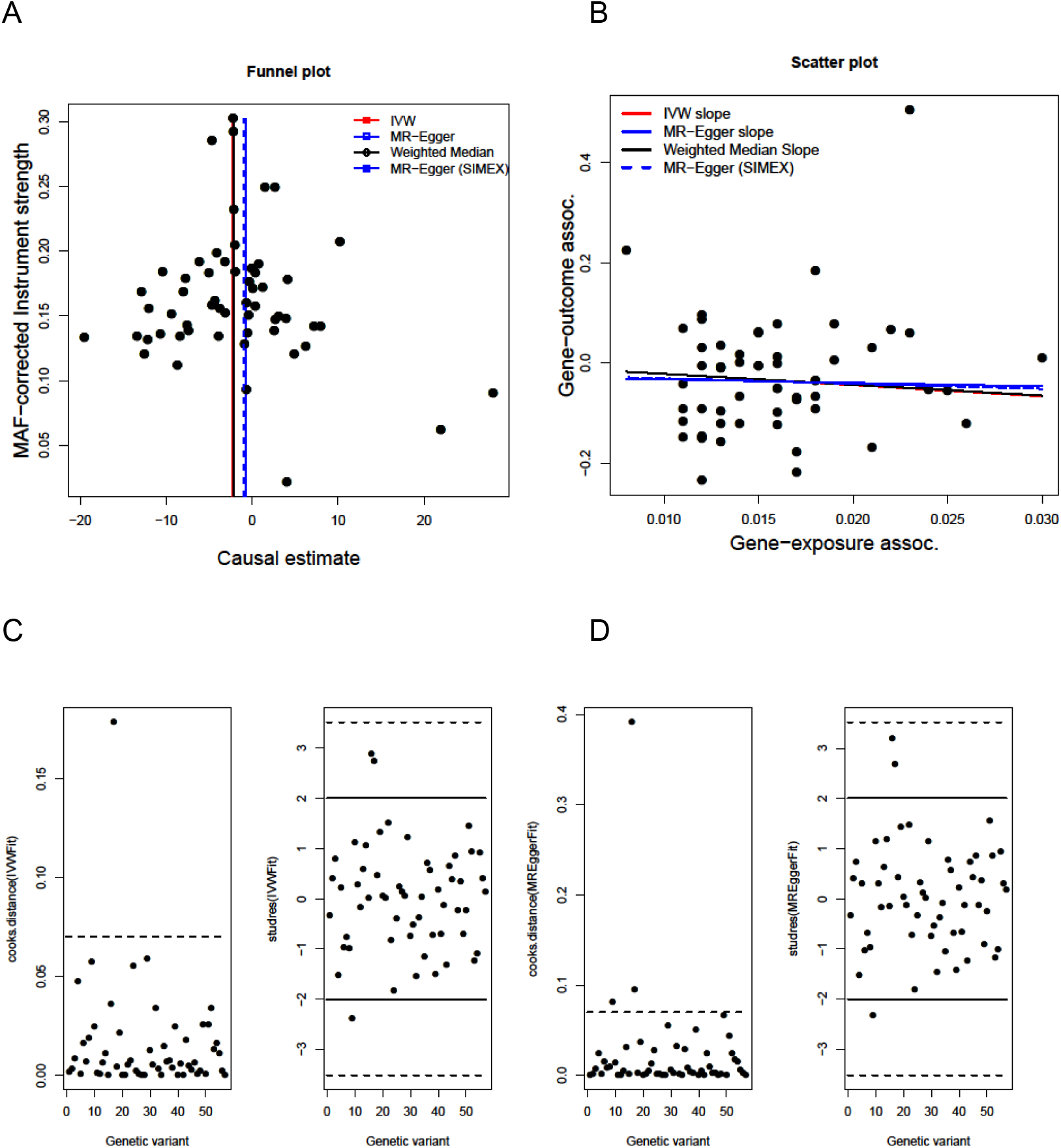
Genetic associations of education with cigarettes per day. Analysis of 38,181 daily smokers (SNP N=57 of a possible 74). A) Funnel plot showing minor allele frequency and causal estimate correlations for each SNP. Coloured lines represent the effect sizes of the different regression analyses. B) Scatter plot showing correlation of genetic associations of education with genetic associations with cigarettes per day. Coloured lines represent the slopes of the different regression analyses. C) Cook’s distances and studentized residuals from inverse variance weighted method. D) as C) from MR Egger method, in order to ascertain the presence of outliers.

There was little evidence of an association between years of education and cotinine levels. However, after assessment of Cook’s distances an outlier SNP (rs113520408) was identified and removed in a sensitivity analysis. This SNP was identified in the initial education GWAS as being worthy of further investigation as it showed sign-discordant effects on height and educational attainment though we could not identify a specific biological rationale for this SNP being an outlier. Inverse variance weighted MR, weighted median MR and MR Egger all produced beta coefficients in the expected direction in this case (i.e., with more years of education being associated with lower cotinine levels), although statistical evidence of association was weak for all methods (e.g., inverse variance weighted beta -0.34, 95% CI -0.67, -0.01). The I^2^_GX_ value was 0.64. The intercept in the MR Egger analysis did not indicate any evidence of pleiotropy (SIMEX corrected beta 0.01, 95% CI -0.02, 0.04). These results are shown in Table 3 and Figure 3.

**Table 3:**
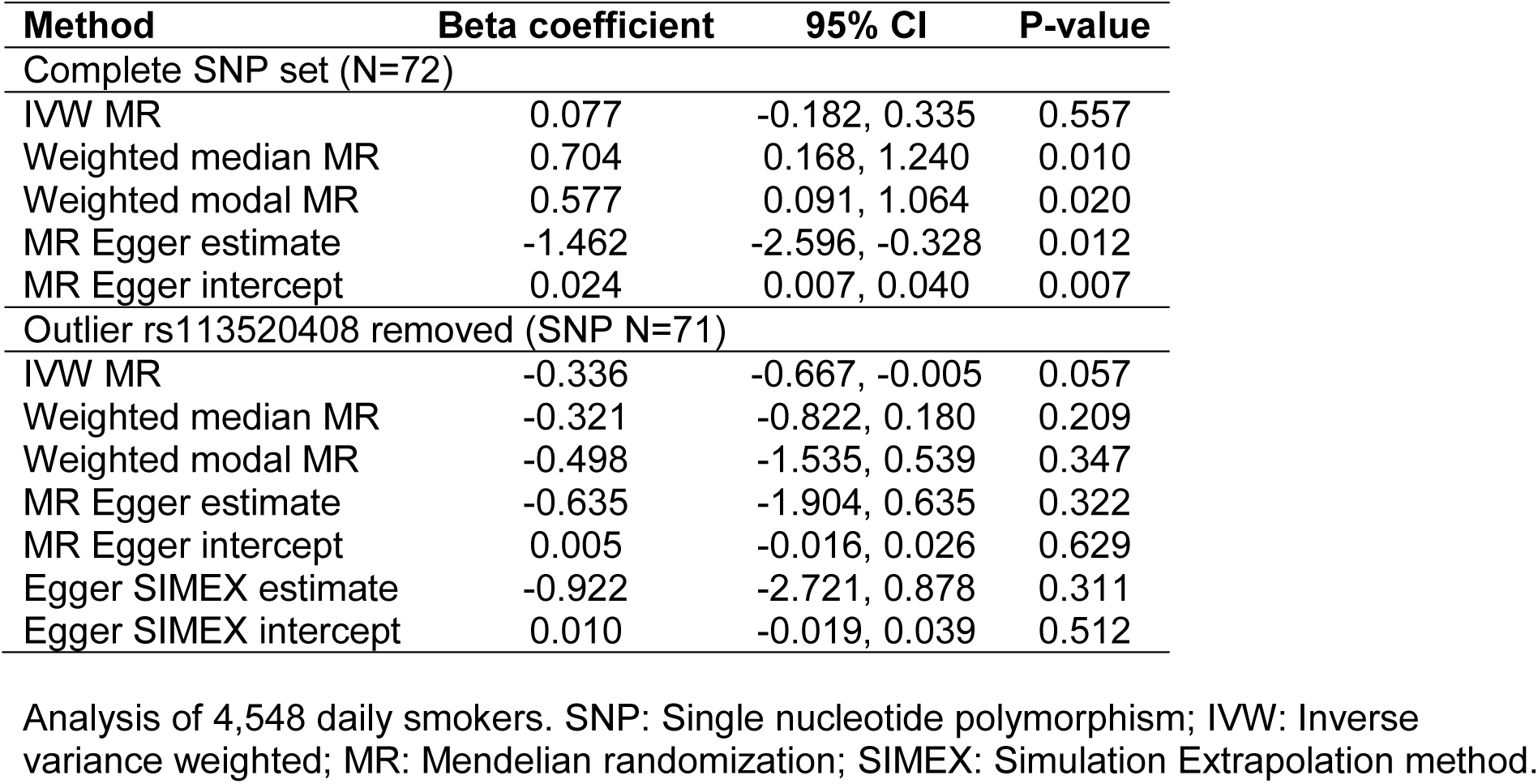
Estimates from various Mendelian randomization methods for the association between education and cotinine levels.

**Figure 3:**
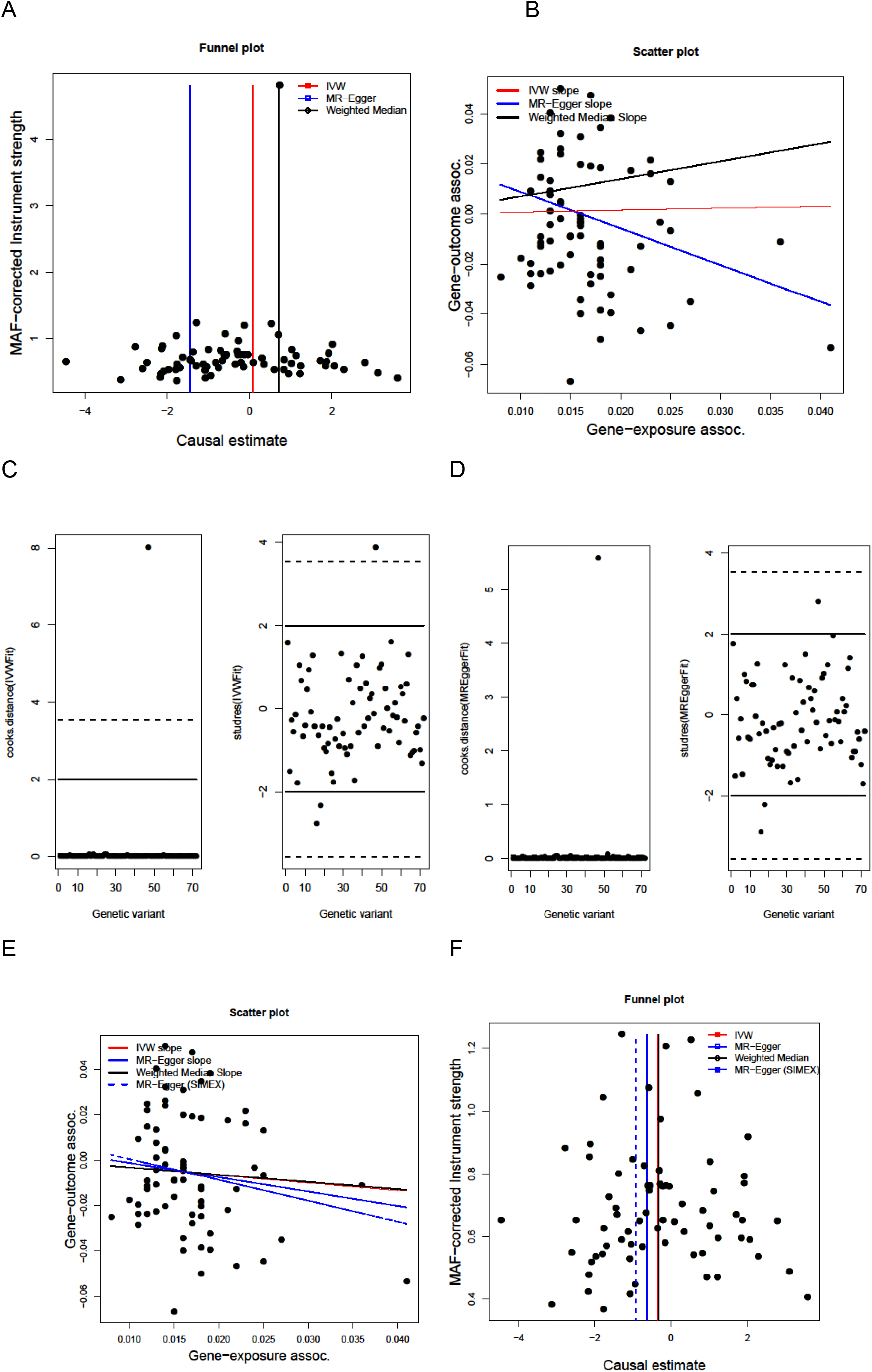
Genetic associations of education with cotinine levels. Analysis of 4,548 daily smokers (SNP N=72 of a possible 74). A) Funnel plot showing minor allele frequency and causal estimate correlations for each SNP. Coloured lines represent the effect sizes of the different regression analyses. B) Scatter plot showing correlation of genetic associations of education with genetic associations of cotinine levels. Coloured lines represent the slopes of the different regression analyses. C) Cook’s distances and studentized residuals from inverse variance weighted method. D) as C) from MR Egger method, in order to ascertain the presence of outliers. E) and F) as A) and B) after excluding outlier SNP rs113520408

### Smoking Cessation

There was strong evidence of an association between more years of association greater likelihood of smoking cessation (beta 0.65, 95% CI 0.35, 0.95). I^2^_GX_ was 0.62 for these analyses. Again, results were similar when using weighted median MR (beta 0.60, 95% CI 0.16, 1.04), weighted modal MR (beta 0.39, 95% CI -0.59, 1.37) and MR Egger (SIMEX corrected beta 0.61, 95% CI -1.14, 2.35), although confidence intervals were wider. The intercept did not indicate any evidence of pleiotropy (SIMEX corrected beta 0.00, 95% CI -0.03, 0.03). These results are shown in Table 4 and Figure 4.

**Table 4:** Estimates from various Mendelian randomization methods for the association between education and smoking cessation.

| Method | Beta coefficient | 95% CI | P-value |
| --- | --- | --- | --- |
| IVW MR | 0.651 | 0.352, 0.950 | $5.54 \times 10^{-5}$ |
| Weighted median MR | 0.600 | 0.160, 1.040 | 0.008 |
| Weighted modal MR | 0.391 | -0.587, 1.369 | 0.433 |
| MR Egger estimate | 0.415 | -0.741, 1.571 | 0.475 |
| MR Egger intercept | 0.004 | -0.015, 0.022 | 0.673 |
| Egger SIMEX estimate | 0.606 | -1.141, 2.353 | 0.500 |
| Egger SIMEX intercept | 0.001 | -0.027, 0.029 | 0.946 |

Analysis of 41,278 individuals (SNP N=57 out of a possible 74). SNP: Single nucleotide polymorphism; IVW: Inverse variance weighted; MR: Mendelian randomization; SIMEX: Simulation Extrapolation method.

**Figure 4:**
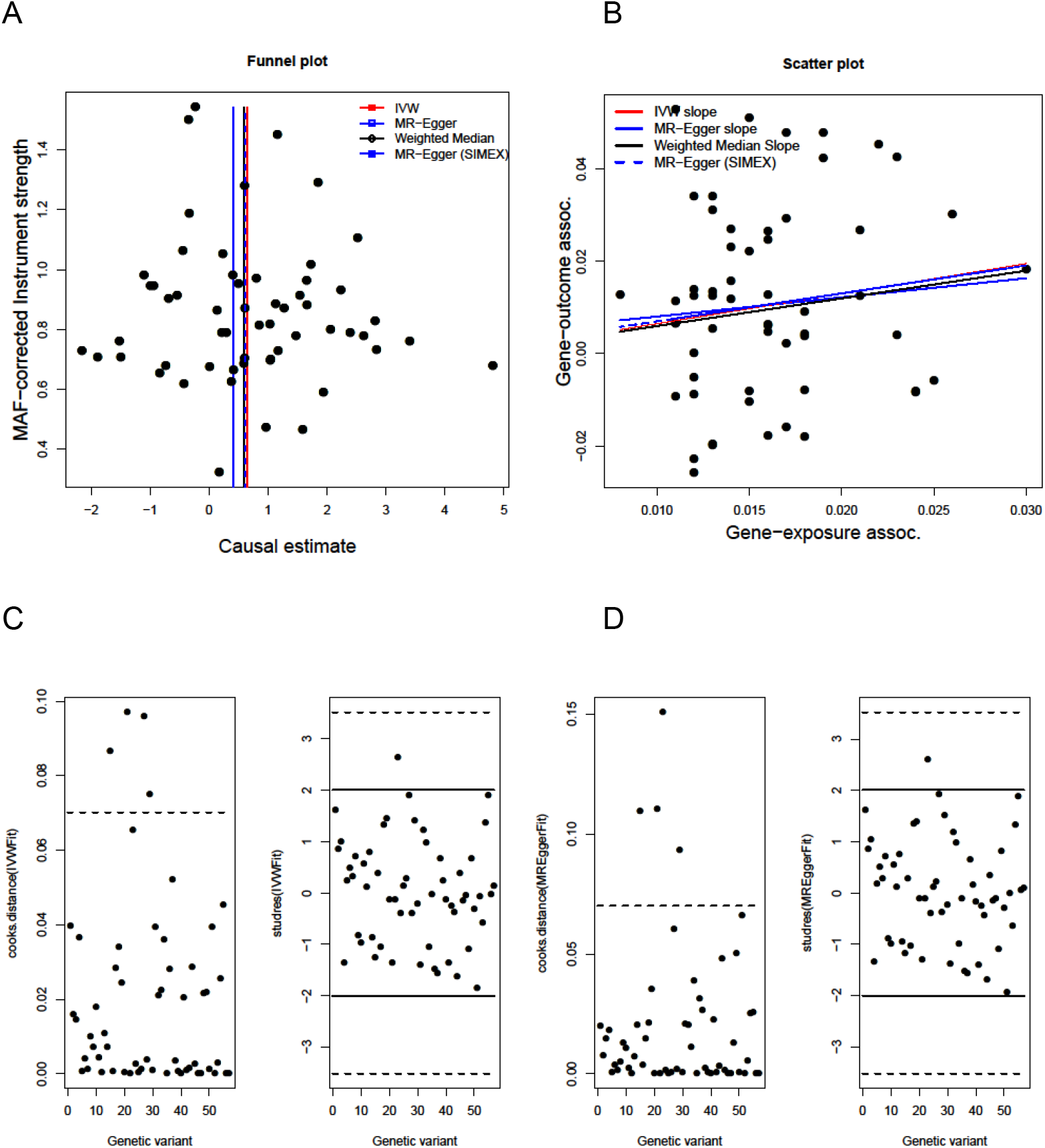
Genetic associations of education with likelihood for smoking cessation. Analysis of 41,278 individuals (SNP N=57 of a possible 74). A) Funnel plot showing minor allele frequency and causal estimate correlations for each SNP. Coloured lines represent the effect sizes of the different regression analyses. B) Scatter plot showing correlation of genetic associations of education with genetic associations with smoking initiation. Coloured lines represent the slopes of the different regression analyses. C) Cook’s distances and studentized residuals from inverse variance weighted method. D) As C) from MR Egger method, in order to ascertain the presence of outliers.

## Discussion

By triangulating evidence from three complementary MR methods that rely on different underlying assumptions, we find evidence that more years in education leads to reduced likelihood of smoking initiation, reduced heaviness of smoking among smokers, and greater likelihood of smoking cessation among smokers. Although statistical evidence was generally weaker when using weighted median and MR Egger methods, results across the methods were broadly consistent in terms of the direction and strength of association observed. Moreover, MR Egger indicated only weak or little evidence of biological pleiotropy. Biological pleiotropy, where one variant has a direct effect on two or more phenotypes, differs from mediated pleiotropy, where a variant impacts on a phenotype via another phenotype. MR assumptions are violated by biological pleiotropy, but not by mediated pleiotropy.

The directionally opposite results obtained using MR Egger for smoking initiation (compared with those obtained using the inverse variance weighted and weighted median methods) is surprising. This may be because smoking initiation might not be a less precise phenotype as some of the others. For example, many more people will try cigarettes than will become daily smokers, and the variants identified in the initiation GWAS could conceivably be measuring another phenotype, such as impulsivity or novelty seeking. MR Egger suggested weak evidence of pleiotropy for this analysis. The results of the cotinine analysis were initially inconsistent, but this seemed to be largely due to the influence of one SNP that was identified as an outlier. Once removed, the estimates from the various MR methods were much more similar, and consistent with the results for the analysis of selfreported cigarettes per day (despite being based on a much smaller sample size).

Our use of multiple smoking phenotypes and various different MR techniques is an important strength. However there are limitations to our results that are important to consider. In particular, not all the genome-wide significant SNPs that predicted educational attainment were available in the outcome GWAS we used, meaning we are not necessarily capturing the full variance with the included variants. Although we were able to identify some proxies, for the analyses that used the TAG consortium we were still missing 17 SNPs.

Our findings may explain the observational associations between educational attainment and adverse health outcomes such as risk of coronary heart disease. Indeed, a recent comment piece argues precisely this, that social rank has an impact on health both on lifestyle behaviours (such as smoking) and via other pathways (19). A recent study concluded that smoking was only a partial mediator of the association between intelligence and mortality, although the measure of smoking used was crude and therefore residual confounding is still possible (20). Our results indicate that education could represent a worthwhile target for intervention. A recent natural experiment exploiting the raising of school leaving age in UK changes found evidence of causal associations of increased schooling on a variety of health and socio-economic factors. While they found little evidence that the one-year increase in schooling was associated with smoking, they did find evidence of a causal association between education and decreased risk of various health outcomes including diabetes and stroke. This inconsistency between these findings and the results we report here could be due to the use of UK Biobank data by Davies and colleagues. Higher educational attainment may be associated with greater likelihood of participation in UK Biobank, while smoking may be associated with lower likelihood of participation. This could lead to collider bias and a consequent attenuation to the null of associations between educational attainment and smoking(21). Policymakers should consider the length and quality of education provision, given growing evidence that it can causally impact on health and health-related behaviours.

## Acknowledgements

This work was supported by the Medical Research Council and the University of Bristol (MC_UU_12013/1, MC_UU_12013/6). JB is supported by an MRC Methodology Research Fellowship (grant MR/N501906/1) MRM and SHG are member of the UK Centre for Tobacco and Alcohol Studies, a UKCRC Public Health Research: Centre of Excellence. Funding from British Heart Foundation, Cancer Research UK, Economic and Social Research Council, Medical Research Council, and the National Institute for Health Research, under the auspices of the UK Clinical Research Collaboration, is gratefully acknowledged.

